# Effect of pH on the thermostability and redox properties of cytochrome *c*_552_ from *Wolinella succinogenes*

**DOI:** 10.1101/2024.03.09.584216

**Authors:** Vitor H. Mordido, Marta S. P. Carepo, Cristina M. Cordas, Navendu Paul, Jörg Simon, Isabel Moura, Sofia R. Pauleta

## Abstract

Cytochrome *c*_552_ from *Wolinella succinogenes* is one of the few examples of a low reduction potential class I *c*-type cytochrome with a mixture of high/low spin state populations observed in its visible spectrum. Analysis of its structural model suggests that the heme is Met/His coordinated and highly solvent-exposed. This supports the hypothesis that it is the solvent accessibility of the propionate groups that controls the reduction potential of small *c*-type cytochromes. The visible spectra obtained at different pH values reveal the presence of a protonable group with a p*K*_a_ of 7.3, which also influences the reduction potential of this small cytochrome *c*_552_ (E_m_^0’^ of 97 ± 5 mV, pH 7.0) and can be either an H_2_O/OH^-^ group distantly coordinating the heme iron, or one of the propionate groups. The thermostability of cytochrome *c*_552_ has been studied by circular dichroism and differential scanning calorimetry, indicating a highly stable protein at pH 5-7 (90 °C to 77 °C).

## 1 Introduction

Denitrifying organisms reduce nitrate to nitrogen in four steps, each catalyzed by a metalloenzyme: nitrate reductase (Sparacino-Watkins et al., 2014), nitrite reductase (Besson et al., 2022), nitric oxide reductase (Shiro, 2012), and nitrous oxide reductase (Pauleta et al., 2013). This enzyme catalyzes the final step of this pathway, denitrification, which is the reduction of nitrous oxide to dinitrogen, according to the Equation 1 (Pauleta et al., 2019).

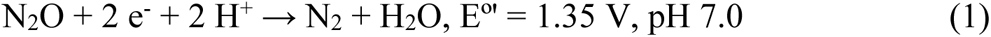

Nitrous oxide reductase is a multiple copper enzyme containing two copper centers: CuA a binuclear electron transfer center and the catalytic site CuZ, which contains four copper atoms. The two electrons required for the reduction are delivered to the CuA center by small redox proteins, T1 copper proteins or *c*-type cytochromes (Dell’Acqua et al., 2008; Fujita et al., 2012).

These small *c*-type cytochromes belong to the class I because they have about 120 residues, with the heme-binding motif near the N-terminus, and the sixth ligand, a methionine residue, 40 residues toward the C-terminus. Thus, the heme is covalently bound to the polypeptide chain by two thioether bonds and it is Met/His hexacoordinated, being usually in a low-spin configuration with a reduction potential of about + 250 mV *versus* NHE at pH 7.0 (Senn and Wüthrich, 1985; Ferri et al., 1996). Structurally these cytochromes have 3-6 α-helices, with the heme being shielded from the solvent (Bertini et al., 2006).

The bacterium *Wolinella succinogenes* can reduce nitrate to ammonia in a respiratory process but is still capable of reducing nitric oxide and nitrous oxide gases (Payne et al., 1982; Kern and Simon, 2009). *W. succinogenes* cytochrome *c* nitrous oxide reductase has been isolated and a small cytochrome *c* of 81 residues has been proposed as its possible physiological electron donor (Teraguchi and Hollocher, 1989; Zhang and Hollocher, 1993). In addition, this small cytochrome *c* is also predicted to mediate electron transport between the Rieske cytochrome *bc* complex and the cytochrome *cbb*_3_ oxidase and was first isolated by our group in 1988 (Moura et al., 1988; Baar et al., 2003; Kern et al., 2010). Visible spectroscopy, as well as NMR studies suggested the presence of an equilibrium between two ligand arrangements around the heme (a high-spin form coexisting with a low-spin form of the heme, with methionine coordination) and a reduction potential lower than usual, +105 m, pH 7.6 (Moura et al., 1988).

Another small *c*-type cytochrome has been isolated in our group from a denitrifying marine bacterium, *Marinobacter nauticus*, cytochrome *c*_552_. Although, also smaller than the usual class I cytochromes, its reduction potential is higher, + 250 mV at pH 7.6 (Saraiva et al., 1994). This small *c*-type cytochrome is always a dimer in solution and no high-spin form was observed. The structural similarities and different spectroscopic properties of these two rather small *c*-type cytochromes are discussed here.

In this paper we describe the heterologous production of cytochrome *c*_552_ from *W. succinogenes*. The spectroscopic characterization of this small *c*-type cytochrome is further investigated by various techniques, correlated with our previous results, and interpreted in the light of the structural model reported here. The thermostability of this protein was studied by differential scanning calorimetry and circular dichroism. The system for its heterologous production will allow us to proceed with electron transfer studies with cytochrome *c* nitrous oxidase reductase.

## 2 Materials and Methods

### 2.1 Chemicals

Unless otherwise stated, all reagents were of analytical or higher grade and were purchased from Sigma-Aldrich, Merck and Fluka. Solutions were prepared in bi-distilled water or Milli-Q water where indicated.

### 2.1 Bioinformatic analysis

The DNA sequence of cytochrome *c*_552_ from *W. succinogenes* (WS0700) was obtained from the GenBank (http://www.ncbi.nlm.nih.gov/genbank) to design the primers. All the analyzed protein sequences were obtained from the Protein database at NCBI (http://www.ncbi.nlm.nih.gov/protein). Multi-sequence alignments were performed using the Clustal Omega (Sievers et al., 2011). Prediction of a signal peptide was performed using SignalP - 6.0 (Teufel et al., 2022). A structural model of the globular domain of *Ws* cytochrome *c*_552_ was created using AlphaFold colab (Jumper et al., 2021), and the heme group was added manually.

### 2.2 Heterologous production of *W. succinogenes* cytochrome *c*_552_

The gene encoding *W. succinogenes* cytochrome *c*_552_ (WS0700), hereafter referred to as *Ws* cytochrome *c*_552_, was amplified by PCR using a set of primers (forward: CATGCCATGGCTGATGGTGCAACCCTCTA, and reverse: CTGGCTCGAGTTACTTGAGCGTGGAGATGTAT) with a NcoI and a XhoI restriction site at the 5’ and 3’-ends, respectively. The amplified fragment encoding the mature cytochrome *c*_552_, excluding its signal peptide (residue 1 to 17, Figure S1, in Supplementary Material), was cloned into a pET22b (+) plasmid, hereafter designated as pET22-*Ws*cytc. The plasmid confers ampicillin resistance and adds an N-terminal signal peptide (*pelB*) to direct the protein to the periplasm. This cloning strategy introduced an additional Met residue, at the N-terminus after cleavage of the signal peptide, giving the mature protein 82 residues.

*Escherichia coli* DH5α (Invitrogen) was used for plasmid propagation. *Ws* cytochrome *c*_552_ was produced in *E. coli* C41(DE3) (Merck) co-transformed with pET22-*Wscytc* and pEC86 (which harbors the *ccm* genes for production of the machinery for *c*-type heme biosynthesis and maturation (Arslan et al., 1998), and confers chloramphenicol resistance). Four to five colonies of the co-transformed *E. coli* C41(DE3) were used to inoculate 50 mL of Luria-Bertani (LB) medium (10 g tryptone, 10 g NaCl and 5 g yeast extract, per liter) supplemented with 100 µg/mL ampicillin and 30 µg/mL chloramphenicol and grown overnight at 37 °C, 210 rpm. Fresh 2xYT medium (16 g tryptone, 5 g NaCl and 10 g yeast extract, per liter), supplemented with the same antibiotics, was inoculated with 1 % of the pre-inoculum. Cultures were incubated under orbital shaking at 37 °C, 210 rpm until an OD_600 nm_ of 0.6 was reached. At this point, gene expression was induced with 0.25 mM IPTG for 18 hours at 30 °C, 120 rpm. Cells were harvested at 8000 ×*g*, 6 °C, 15 min, and resuspended in 50 mM Tris-HCl, pH 7.6 containing protease inhibitors (cOmplete™, Mini, EDTA-free, Protease Inhibitor Cocktail Tablets, Roche).

### 2.3 Purification of heterologous *W. succinogenes* cytochrome *c*_552_

The periplasmic fraction was obtained by 4 freeze-thaw cycles and separated from spheroplasts and cell debris by centrifugation at 39000 ×*g*, 6 °C, 45 min. Purification was performed in a single chromatographic step, using a cationic exchange chromatography. The periplasmic fraction was diluted 10× with cold milli-Q water and loaded onto a CM52 (Cytiva) cation-exchange chromatographic column (12.5 cm x 3 cm Ø, 90 mL), equilibrated with 5 mM sodium phosphate buffer, pH 7. Protein not adsorbed on the matrix was eluted with 5 mM sodium phosphate buffer, pH 7, and then a linear gradient between 0 and 500 mM NaCl was applied. The *Ws* cytochrome *c*_552_ eluted with 125 mM NaCl, and the fractions with A_408.5nm_/A_280nm_ above 4.7 were considered pure, combined, and concentrated over a 3 kDa MWCO membrane using a Vivaspin 15 centrifugal device (Sartorius). The final *Ws* cytochrome *c*_552_ fraction was buffer exchanged to 5 mM Na-phosphate pH 7.0 using a desalting PD-10 column (Cytiva). A 15 % SDS-PAGE and 10 % PAGE stained for protein (Coomassie blue) and heme content (Goodhew et al., 1986) was also used throughout the purification to verify the protein purity. *Ws* cytochrome *c*_552_ was stored in small aliquots at - 80 °C until further use. Protein and heme content was estimated in the same solution for which the CD spectra in the far-UV region was acquired: protein was quantified using the extinction coefficient at 205 nm determined for CD (see below) and heme content by the pyridine hemochrome assay (Berry and Trumpower, 1987).

### 2.4 Spectroscopic characterization

The UV-visible spectra were recorded on a Shimadzu UV-1800 spectrophotometer, connected to a computer, using a 1 cm path quartz cuvette. The concentration of the protein was estimated using the reported molar extinction coefficient of 80.1 mM^−1^cm^−1^ at pH 7.6, 100 mM K-phosphate buffer (Moura et al., 1988). *Ws* cytochrome *c*_552_ in the reduced state was obtained by reduction with a solution of sodium dithionite, with a final concentration of 5 mM.

Circular dichroism spectra were acquired in an Applied Photophysics ChirascanTM qCD spectrometer (Leatherhead, Surrey, UK). Far-UV (190–260 nm) spectra were acquired in a sample of 8.5-10.6 μM *Ws* cytochrome *c*_552_, in 5 mM Na-phosphate buffer, pH 5.0, 6.0, 7.0, 8.0, and 9.0, using a 1 mm path length cuvette with a total volume of 300 μL. In the visible region (260–800 nm) the spectra were recorded for a 35 μM *Ws* cytochrome *c*_552_ sample, in 5 mM K-phosphate buffer, pH 7.0, using a 10 mm path length cuvette with a total volume of 2.3 mL. The concentration of the samples was determined using the absorbance at 205 nm and the extinction coefficient of 290700 mM^−1^•cm^−1^, based on the amino acid content (Anthis and Clore, 2013), or the absorbance at 408 nm and the extinction coefficient mentioned before. CD data were reported in mean residue ellipticity ([θ]_MRE_) as a function of wavelength. The [θ]_MRE_ is the CD raw data (in deg) corrected for the *Ws* cytochrome *c*_552_ concentration of the solution using Equation (1):

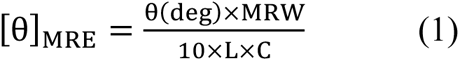

where MRW (mean residue weight) is the molecular mass of *Ws* cytochrome *c_552_* divided by the number of peptide bonds (81), C is the concentration of the protein in the sample in g•mL^−1^, and L is the pathlength of the cell in cm.

The CD spectra between 190 and 260 nm are an average of three spectral acquisitions at 25 °C, with a bandwidth and step-size of 1 nm, and acquired with a time per point of 3 s. The far-UV data analysis to determine the secondary structure content was performed using the BeStSel server (https://bestsel.elte.hu/) (Micsonai et al., 2021; Micsonai et al., 2022). The CD spectra between 260 and 800 nm are an average of three spectral acquisitions at 25 °C, with a bandwidth and step-size of 1 nm, and acquired with a time per point of 3 s. The temperature-dependent CD spectra were acquired between 10 °C and 94 °C, with a stepped ramp mode of 1 s per point, an increase of 2 °C for each measurement, and a stabilization period of 1 min between each point.

The unfolding process was analyzed using the Gibbs–Helmholtz method - to fit the change in ellipticity at a single wavelength as a function of temperature, considering a two-state transition from a folded native state to an unfolded state, and assuming that the heat capacity of the folded and the unfolded states are equal, ΔCp = 0 (Greenfield, 2004). For the Gibbs–Helmholtz method, the experimental data were fitted using Equations (2)-(5) and using the SOLVER add-in program of Microsoft Excel. The molar ellipticity at any given temperature (T), [θ]_T_, is given by Equation (2):

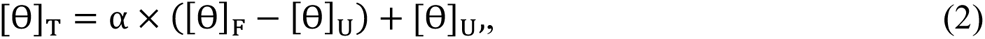

in which the fraction folded at any given temperature, α, is given by Equation (3):

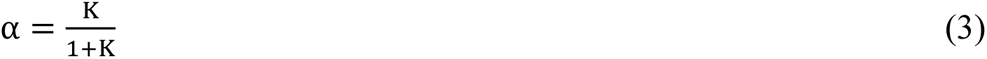

and the folding constant, K, at any given temperature is given by Equation (4):

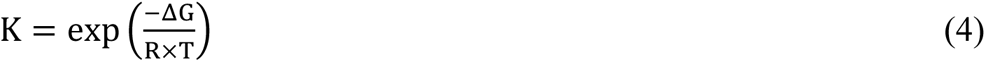

and ΔG is given by Equation (5):

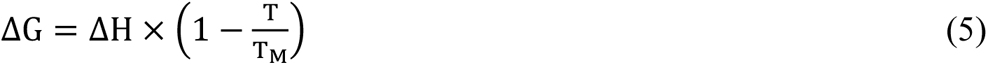

in which T_M_ is the temperature at which the fraction folded, α, is 0.5.

### 2.5 Differential Scanning Calorimetry

For the differential scanning calorimetry (DSC) experiments, the NanoDSC instrument (TA Instruments) was loaded with degassed buffers (baselines and reference cell) and protein solution (83-93 µM *Ws* cytochrome *c*_552_ in the sample cell). Each protein sample was passed through a desalting PD-10 column (Cytiva) equilibrated in the appropriate buffer (5 mM K-phosphate buffer at pH 5.0, 6.0, 7.0, 8.0, and 9.0) and then diluted to the desired concentration in the same buffer. The temperature was raised from 10 to 120 °C at a scan rate of 1 °C/min, and 6 bar. The thermograms were analyzed with TA instruments NanoAnalyze software using a non-two-state model to fit the data and obtain the melting temperature (*T_M_*), the calorimetric (*ΔH*) and van’t Hoff (*ΔH_v_*) enthalpies. The corresponding baseline was subtracted from each sample scan.

### 2.6 Determination of the reduction potential

Cyclic voltammetry assays were performed in a single compartment cell with a 3-electrode configuration. A cysteamine-modified gold disk (ϕ = 2 mm) was used as the working electrode and a platinum foil and a saturated calomel electrode (SCE) were used as the secondary and reference electrodes, respectively. The gold electrode was modified by immersion in a 2 mM cysteamine solution for 30 min at room temperature (RT), followed by washing by immersion in deionized water. The protein (5 μL of approx. 550 μM) was then placed on the modified gold electrode and a cellulose membrane (3.5 kDa cut-off) was then applied for protein entrapment and studied in a thin-layer regime. The used electrolyte was a mixed 20 mM phosphate/acetate buffer / 0.1 M NaCl, at different pH values or only 20 mM phosphate / 0.1 M NaCl, pH 7. Assays were performed at RT in a strictly anaerobic environment inside an anaerobic chamber (c_O2_<0.1 ppm). Different potential windows and scan rates were tried. All the potentials were converted and presented in the normal hydrogen electrode (NHE) reference scale. Controls were obtained using the same procedure and conditions without the presence of the protein.

## 3 Results

### 3.1 Primary Sequence Analysis and Structural Model

Analysis of the primary sequence of the mature cytochrome *c*_552_ from *W. succinogenes* (81 residues) reveals the presence of an N-terminal heme-binding motif and a conserved methionine residue, about 40 residues away towards the C-terminal, as distal axial ligand of the heme iron. Such characteristics indicate that this *c*-type cytochrome belongs to class I. The alignment with other small *c*-type cytochromes (Figure 1), one of which, *M. nauticus* cytochrome *c*_552_, is also an electron donor of a nitrous oxide reductase, shows these conserved features.

**Figure 1.**
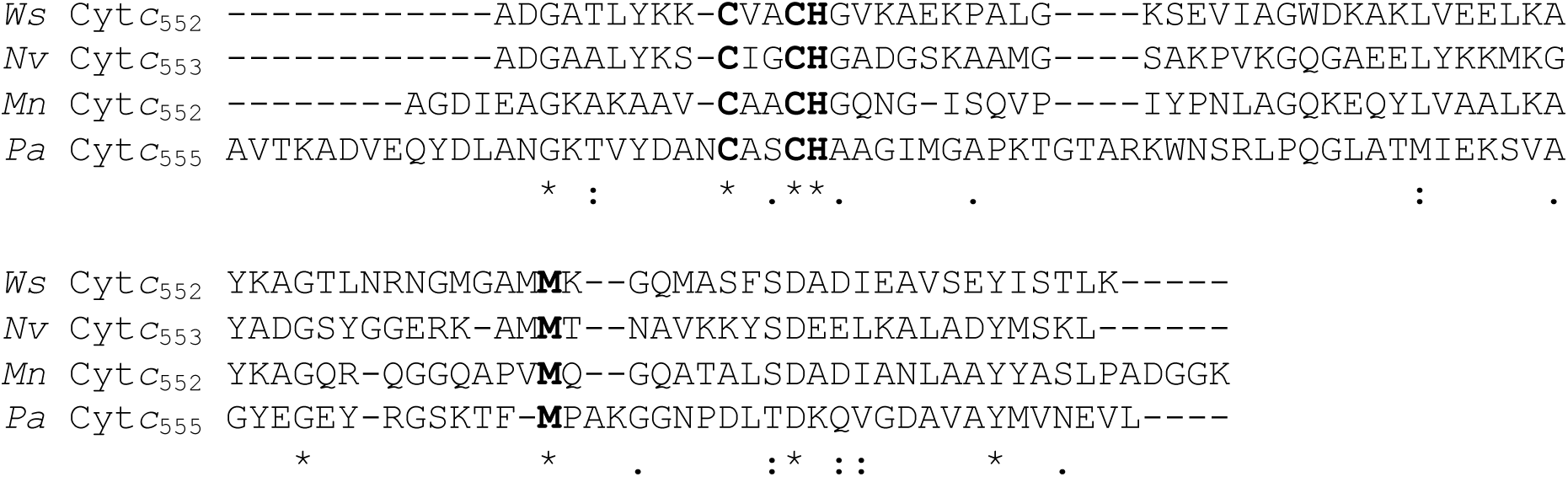
Primary sequence alignment of the mature cytochrome *c*_552_ from *W. succinogenes* with other small cytochromes *c*, with similar redox properties and structure. The heme binding motif and the axial ligands are shown in bold. Asterisks, colons or stops below the sequence indicate identity, high conservation or conservation of the amino acids, respectively. Legend: *Ws* Cyt*c*_552_ (*Wolinella succinogenes* cytochrome *c*_552_, Q7MS72), *Nv* Cyt*c*_553_ (*Nitratidesulfovibrio vulgaris* Hildenborough cytochrome *c*_553_, P04032), *Mn* Cyt*c*_552_ (*Marinobacter nauticus* cytochrome *c*_552_, P82903) and *Pa* Cyt*c*_555_ (*Prosthecochloris aestuarii* cytochrome *c*_555_, P00124).

The structural model of this protein was obtained using AlphaFold colab (Figure S2, in Supplementary Material), and the heme group was manually added so that the imidazole group of His15 would axially coordinate the heme iron, and the sulfur atoms of the two cysteine residues would be at a distance to form a thioether bond with the vinyl groups at the β-pyrrole positions 2 and 4 of the heme (Figure 2A). This structure shows that it is composed by 4 α-helices, and that the heme group is more solvent exposed than in the case of *Paracoccus denitrificans* cytochrome *c*_550_ (Figure 2D), since the loops covering the heme and 1 extra helix are missing. The comparison of its structure with that of two other small *c*-type cytochromes (Figure 2), those isolated from *M. nauticus* and *Nitratidesulfovibrio vulgaris* Hildenborough, formerly known as *Desulfovibrio vulgaris* Hildenborough, shows that their structure and heme accessibility are similar, although in the case of *M. nauticus* the heme is slightly less exposed to the solvent due to its dimeric form (Figure S9, in Supplementary Materials).

**Figure 2.**
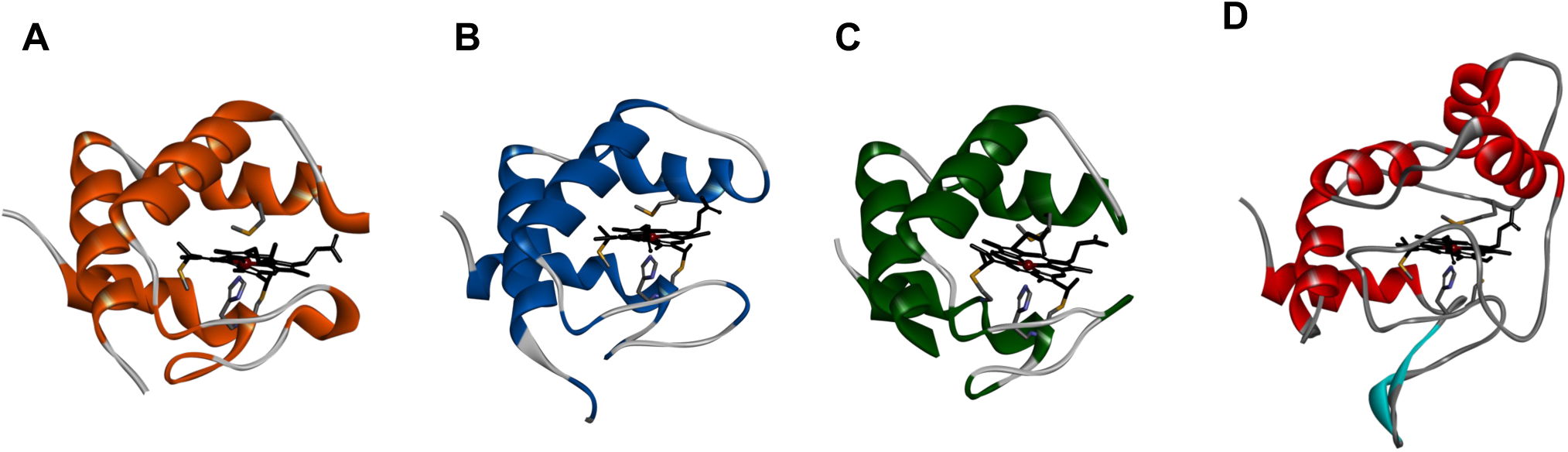
Structure of small *c*-type cytochromes. Comparison of the structural model of cytochrome *c*_552_ from *W. succinogenes* obtained using the Alphafold colab Panel (A), with the structures of *N. vulgaris* Hildenborough cytochrome *c*_553_ (PDB ID 1DVH) (Panel B), *M. nauticus* cytochrome *c*_552_ (PDB ID 1CNO) (Panel C, showing only the monomer) and *P. denitrificans* cytochrome *c*_550_ (PDB ID 1COT) (Panel D). The heme group and axial ligands are shown as sticks colored by atom. Figure was prepared in BIOVIA Discovery Visualizer Studio using the coordinates indicated above.

### 3.2 Heterologous production of *Wolinella succcinogenes* cytochrome *c*_552_

The DNA fragment amplified by PCR and inserted into the pET22b expression vector encoded only the globular region of *Ws* cytochrome *c*, starting at A18, and due to cloning an additional Met residue was added at the N-terminus (Figure S1, in Supplementary Material). This recombinant protein has 82 residues, after removal of the PelB signal peptide, and an expected molecular mass of 9342.7 Da (8728.2 Da plus 614.5 Da, which is the sum of the molecular mass of the polypeptide chain and one heme group).

The *Ws* cytochrome *c* was isolated from the periplasm of *E. coli* in a single chromatographic step, consisting of a cationic exchange chromatography, since the pI of this protein is 8.6. The cytochrome *c* was considered pure according to its SDS-PAGE and PAGE profile (Figure S3, in Supplementary Material) with a single band, and this fraction had a purity ratio (A_552nm_ – A_570nm_ [reduced form])/A_280nm_ [oxidized form]) of 1.20. This procedure has an average yield of 46 mg of pure *Ws* cytochrome *c* per liter of growth medium. The heme/protein ratio of the purified cytochrome *c*, as determined by heme and protein concentration (see Materials and Methods), was 0.89 ± 0.02, confirming the presence of 1 *c*-type heme bound to the polypeptide chain, as expected.

### 3.3 Spectroscopic Characterization

The UV-visible spectrum of the heterologous cytochrome *c* shows similar spectral features to the same protein isolated from *W. succinogenes* (Moura et al., 1988) (Figure 3A), namely the Soret band at 409 nm, and absorption bands with maxima at 620 nm and 695 nm, indicating the presence of a high-spin state, and a methionyl residue (Met59, numbering according to the mature sequence), respectively. The extinction coefficient estimated based on the heme content is identical to that reported, 80.1 mM^−1^cm^−1^ at pH 7.6 (Moura et al., 1988).

**Figure 3.**
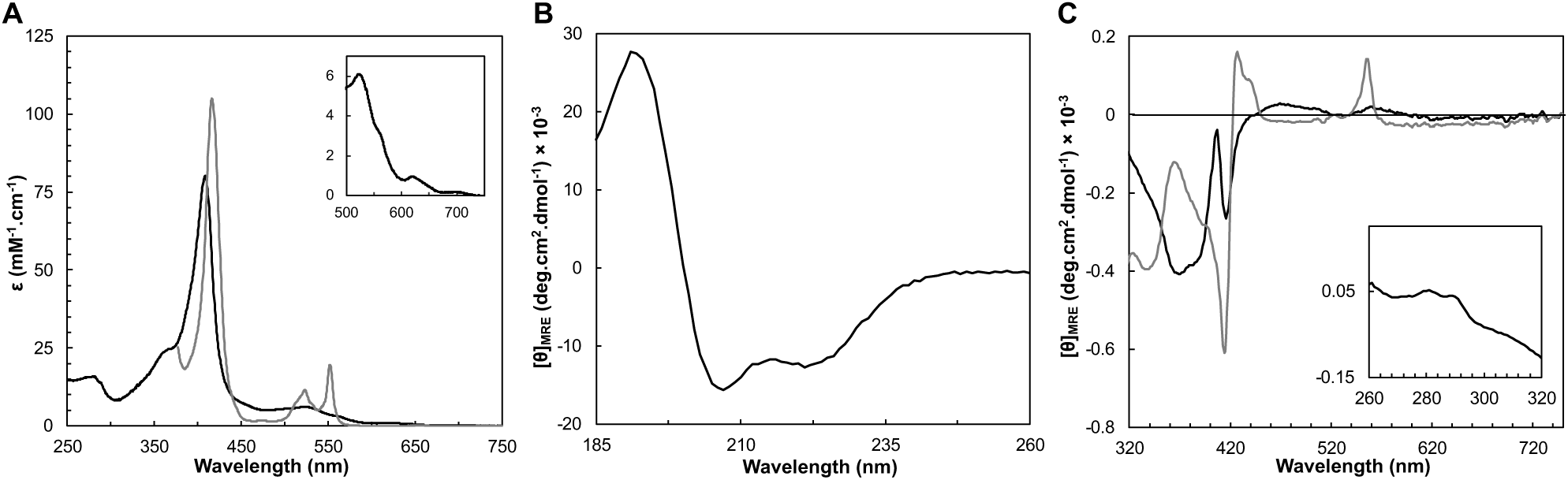
Spectroscopic features of *Ws* cytochrome *c*_552_ in the as-isolated (black line) and reduced state (grey line) at pH 7.0. **(A)** UV-visible spectra acquired in the as-isolated (black line) and reduced state (grey line). Inset shows the Q-band region. Circular dichroism spectra in the far-UV **(B)** and visible **(C)** regions acquired in the two oxidation states, at 25 °C. The inset in panel C shows the near-UV region of the CD spectrum of the oxidized state.

After reduction, the Soret band shifts to 417 nm, with the appearance of the β and α bands at 523 nm and 552 nm, respectively (Figure 3A). The maximum absorption of the α-band at 552 nm, designates this cytochrome as cytochrome *c*_552_, as has been for others (Hon-nami and Oshima, 1977; Saraiva et al., 1994; Samyn et al., 1998).

As previously observed, the high-spin state decreases with increasing pH, as the ratio A_621nm_/A_695nm_ decreases (from pH 5.0 to pH 7.0 the ratio is around 6.0, decreasing to 4.7 and 4.5 at pH 8.0 and 9.0, respectively) (Table 1), indicating a p*K*a around 7.5 (Figure 5B), as previously reported (inflection point at pH 7.3 for the absorbance at 620 nm) (Moura et al., 1988).

**Table 1:** Spectral features and thermostability parameters of *Ws* cytochrome *c*_552_ at different pH values.

|  | pH 5 | pH 6 | pH 7 | pH 8 | pH 9 |
| --- | --- | --- | --- | --- | --- |
| $A_{621nm}/A_{695nm}$ | 5.9 | 5.6 | 5.3 | 4.7 | 4.5 |
| $T_M$ (°C) (CD) | >90 | >85 | >80 | 72.3 ± 0.2 | 72.5 ± 0.1 |
| $T_M$ (°C) (DSC) | 90 ± 0.02 | 83 ± 0.03 | 77 ± 0.02 | 72 ± 0.02 | 70 ± 0.01 |
| $\Delta H_{cal}$ (kJ/mol) | 73 | 101 | 75 | 87 | 98 |
| $\Delta H_{vH}$ (kJ/mol) | 486 ± 3 | 337 ± 2 | 441 ± 3 | 382 ± 2 | 322 ± 1 |
| $\Delta H_{cal}/\Delta H_{vH}$ | 0.2 | 0.3 | 0.2 | 0.2 | 0.3 |
| $\theta_{222nm}/\theta_{208nm}$ | 0.77 | 0.77 | 0.81 | 0.78 | 0.78 |
| α-helix | 28% | 27% | 27% | 32% | 26% |
| β-sheet | 12% | 12% | 10% | 9% | 12% |

The folding state of a protein and its secondary structure content can be determined by analyzing its circular dichroism spectra in the far-UV region (190-260 nm). The CD spectrum of the as-isolated *Ws* cytochrome *c*_552_ shows a positive peak at 192 nm, and two negative peaks at 208 and 222 nm, with a ratio θ_222nm_/θ_208nm_ of 0.8. These features indicate the presence of a folded protein composed mainly of α-helix (Figure 3B). This spectrum is similar to the one of cytochrome *c*_552_ from *Hydrogenobacter thermophilus* and cytochrome *c*_553_ from *N. vulgaris* Hildenborough, cytochromes *c* composed mainly of α-helix (Wittung-Stafshede, 1999; Oikawa et al., 2005). The secondary structure content was estimated by analyzing the far-UV spectrum at 25 °C, pH 7.0 with the BeStSel algorithm (Micsonai et al., 2021; Micsonai et al., 2022), and indicates the presence of 27 % α-helices, with 10 % of β-sheets (Table 1). These values are in agreement with the analysis of the coordinates of the structural model of this protein that predicts a protein mainly α-helix with a small percentage of structure as β-sheets (40% α-helices, with 7% of β-sheets).

The near-UV and visible CD spectra of the oxidized *Ws* cytochrome *c*_552_ were collected in a pH range of 5-9 (Figure S6, in Supplementary Material). In the near-UV region, the presence of two positive bands at 282 nm and 290 nm is observed, which can be attributed to the single tryptophan residue present in the sequence, W33, which is located near the heme propionates in the predicted structure (Figure S10, in Supplementary Material). These bands do not change over the pH range studied. The visible region of the CD spectrum of the oxidized *Ws* cytochrome *c*_552_ at pH 7 has a broad negative peak between 320 nm and 400 nm with a maximum at 370 nm (Figure 3C), which has also been observed in other *c*-type cytochromes (Vinogradov and Zand, 1968). A negative well-defined S-shaped band is observed with a negative minimum at 407 nm and a negative maximum at 416 nm corresponding to the Soret Cotton effect. This Cotton effect on the Soret band of CD spectra has usually been associated with transitions of the heme with nearby aromatic side chains (Hsu and Woody, 1971; Pielak et al., 1986) and is a fingerprint of heme integrity. The visible CD spectra do not change significantly with the pH (Figure S6, in Supplementary Material, suggesting that the tertiary structure of the protein, and in particular the heme pocked is not being affected by changes in the pH. The fact that the Soret band is negative can also be explained by parallel propionate groups on the porphyrin ring (Nagai et al., 2015), that again remains unchanged with pH for *Ws* cytochrome *c*_552_. Upon reduction, the broad band with a maximum at 370 nm becomes a negative minimum and the Soret CD spectrum shows a well-defined sharp band with an inflection point at 422 nm, and a negative peak at c.a. 417 nm, followed by a positive peak at 433 with a shoulder at 440 nm. There is also a positive peak at 556 nm, which corresponds to the peak of the α-band. The changes observed in the visible CD spectra between the oxidized and the reduced forms indicate that the heme environment changes upon reduction. This has also been observed for other *c-*type cytochromes (Vinogradov and Zand, 1968).

### 3.4 Effect of pH in the Thermostability

The influence of pH on the thermostability of cytochrome *c*_552_ was studied by CD in the far-UV region and differential scanning calorimetry (DSC) (Figure 4), which provide complementary information on the unfolding equilibrium. The CD in the far-UV region monitors the change in secondary structure content, while DSC provides information about the overall unfolding process of the protein.

**Figure 4.**
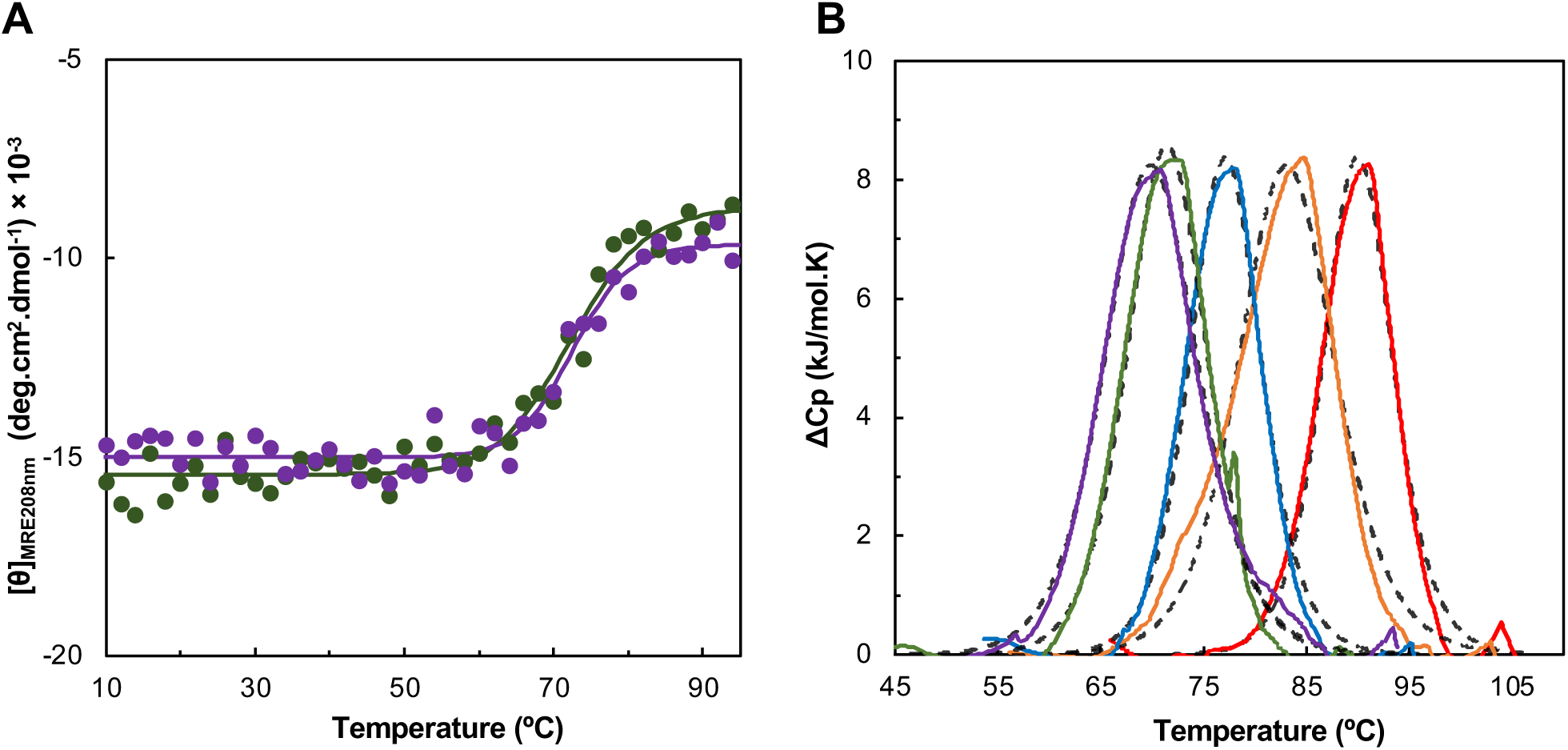
Effect of pH in the thermostability of *Ws* cytochrome *c*_552_ studied by circular dichroism (A) and differential scanning calorimetry (B). Panel A shows the molar ellipticity at 208 nm obtained at different temperature for pH 8.0 (green) and pH 9.0 (purple). The experimental data were fitted using the equations for a two-state transition model (solid line). Panel B shows the thermogram of *Ws* cytochrome *c*_552_ obtained by DSC at pH 5.0 (red line), 6.0 (orange line), 7.0 (blue line), 8.0 (green line) and 9.0 (purple line), each fitted to a one-peak transition model for each case (dashed black line).

**Figure 5.**
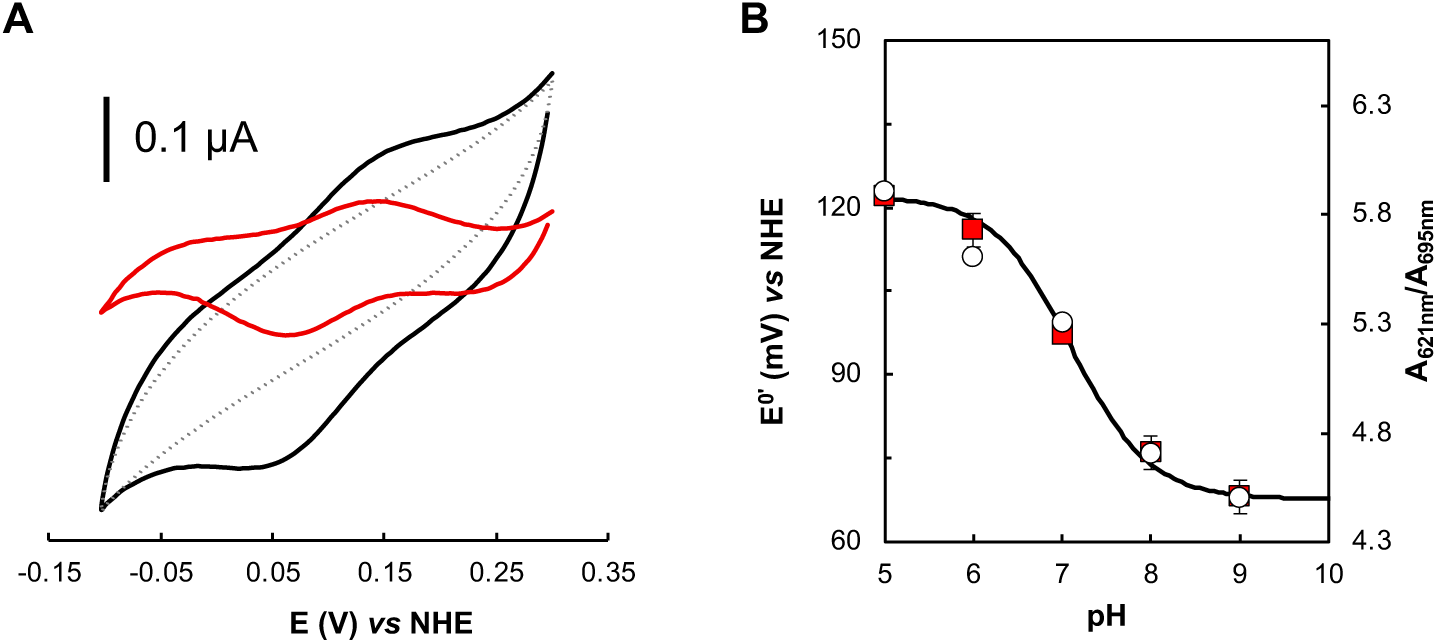
Effect of pH in the reduction potential of *Ws* cytochrome *c*_552_ studied by cyclic voltammetry. (A) Representative voltammogram of the *Ws* cytochrome *c*_552_ (with baseline subtraction, black line, and after control subtraction, red line), and the corresponding control in the absence of the protein (grey line) attained on cysteamine modified gold electrode, at scan rate of 50 mV.s^-1^, 20 mM phosphate buffer / 0.1 M NaCl, pH 7.0, RT, in strict anaerobic environment (anerobic chamber, O_2_ < 0.1 ppm). (B) Formal reduction potential dependence on the pH (red squares). Experimental conditions: average of all replicates (min. 3) and corresponding standard deviation. The data were fitted considering a single protonable group with a p*K*_ox_ and a p*K*_red_, of 6.9 and 7.3, respectively. In the secondary yy axis is represented the A_621nm_/A_695nm_, that shows the same pH profile.

The far-UV CD spectra at different pH values, acquired at 25 °C have similar spectral features to those described at pH 7.0: the peaks have the same maxima and minima (no shifts were observed) (Figure S5, in Supplementary Material). However, the ratio of **θ_222nm_/ θ_208nm_** decreases at higher and lower pH values, indicating that there is a change in the alpha-helix content. Analysis of these spectra using the BeStSel algorithm confirms this small change in the secondary structure content (Table 1), with a decrease in alpha-helix content at pH 5.0, 6.0 and 9.0.

The thermostability studied by CD in the far-UV region shows that this protein is highly stable up to 60 °C, independent of pH (Figure 4A). The T_M_ could only be estimated for pH 8.0 and 9.0, because at lower pH values no stabilization of the ellipticity up to 94 °C was observed. At these two pH values, the estimated T_M_ was 73 °C, but the process is not reversible (Figure S7, in Supplementary Material).

The thermogram obtained by DSC shows a single sharp endothermic peak, which was fitted to a one-peak transition model, from a transition temperature, and a van’t Hoff enthalpy of unfolding, ΔH_vH_, was estimated (Table 1). The shape of the thermogram shows that there is a sharp decrease, indicating that the protein aggregates in solution (Figure S4, in Supplementary Material), which is confirmed by the small ratio ΔH_cal_/ΔH_vH_ (Table 1). This phenomenon occurs for all the pH values studied. The unfolding temperature shows the same trend as that observed for CD, indicating that this cytochrome *c*_552_ is more stable at lower pH values.

### 3.5 Reduction potential at different pH values

The results show that *Ws* cytochrome *c*_552_ has a quasi-reversible electrochemical behavior, as expected considering similar protein systems (Santos et al., 2015; Teixeira et al., 2019). The cyclic voltammogram at pH 7.0 shows a redox process, with broad anodic and cathodic peaks (Figure 5A). The formal reduction potential calculated from the average of E_pa_ and E_pc_ is +97 ± 5 mV *vs* NHE. This experimentally estimated formal reduction potential is in agreement with the previously reported value for this protein of +105 ± 15 mV, obtained by potentiometry (Moura et al., 1988). The relative difference can be explained by the different techniques used.

The study of the redox behavior at different pH values, shows that at pH values lower than pH 7.0, the potential waves are relatively better defined, and a single redox process is observed (Figure S8, in Supplementary Material). Above pH 7.0, a broadening of the waves (both cathodic and anodic) occurs, but it is not possible to distinguish the processes. This behavior, if only pH 7.0 was considered, could be related to some dispersion of the heterogeneous electron transfer rate constants, due to multiple electron transfer pathways. However, the global redox behavior at different pH values points to the coexistence of two slightly different protein forms regarding the heme environment, which is consistent with the hypothesis of having an oxygenated species (OH^-^) or a water molecule binding/leaving at the distal axial position. The pH dependence of the redox potential considering only one acid/base group model is fitted with a p*K*_a_ values for the oxidized and reduced states: p*K*_ox_ and a p*K*_red_ of 6.9 and 7.3, respectively, with a calculated theoretical reduction potential of 0.122 V (pH 0) (Figure 5B).

## 4 Discussion

### 4.1 Structure and redox properties

*W. succinogenes* cytochrome *c*_552_ is an 81-residue *c*-type cytochrome that has been successfully heterologously produced in *E. coli*. It is an interesting protein because it is one of the shortest *c*-type cytochromes isolated to date and has unusual spectroscopic and redox properties, namely a low reduction potential (+97 mV, pH 7.0) and the coexistence of a high and a low-spin forms at room temperature. Other small cytochromes have a similar reduction potential, but the high/low-spin coexistence has not been reported (Table 2). Comparing their model structure, presented here, for the first time, it is clear that due to the shorter polypeptide chain, the heme is more exposed to the solvent, and not in the usual hydrophobic environment with only the exposed heme edge to the solvent (Figure 1). This solvent exposure may explain their lower reduction potential. An exception is the cytochrome *c*_552_ from *M. nauticus*, which has a much higher reduction potential despite being composed of 88 residues and having a similar structure (Table 2). However, this cytochrome *c*_552_ is a dimer in which the propionate groups are shielded from the solvent because they are located at the dimer interface (Figure S9, in Supplementary Material).

**Table 2.**
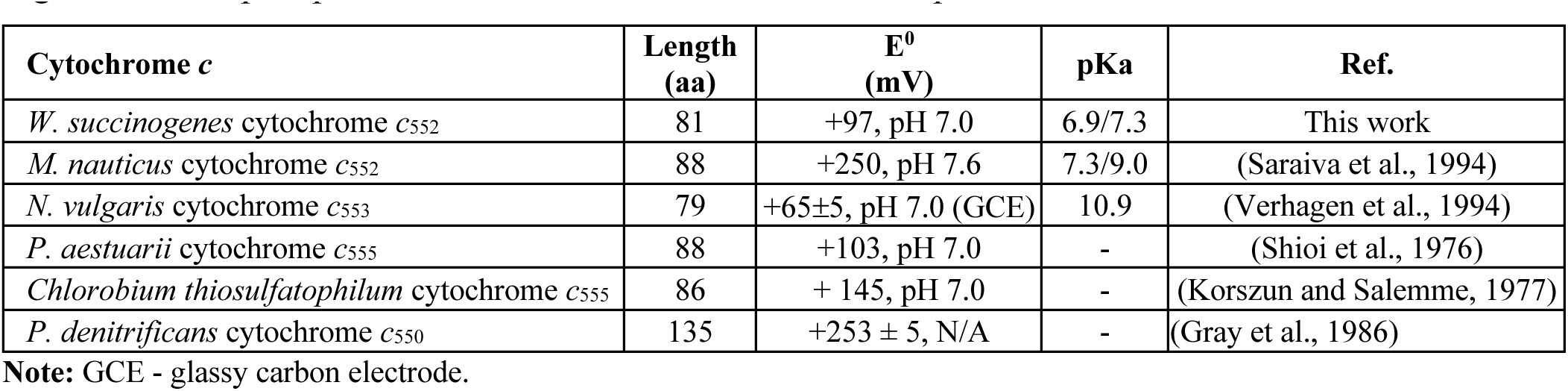
Reduction potential and length of mature small class I *c*-type cytochromes from different organisms. The pKa presented was observed for the reduction potential.

### 4.2 Effect of pH

The effect of pH on the redox properties of *W. succinogenes* cytochrome *c*_552_ differs from that reported for *N. vulgaris* cytochrome *c*_553_, both in the shape of the cyclic voltammograms and in the estimated p*K*a. Similar to other class I *c*-type cytochromes, *N. vulgaris* cytochrome *c*_553_ shows no effect on the reduction potential up to pH 10.0 (Verhagen et al., 1994), after which there is a marked decrease in the reduction potential, which has been attributed to the loss of the methionine as a distal axial ligand with a p*K*_a_ of 10.9 (Verhagen et al., 1994). The pH dependence of *Ws* cytochrome *c*_552_ was studied only up to pH 9.0 to avoid its destabilization close to its pI (of 8.6) and shows a close p*K*_ox_/p*K*_red_, of 6.9/7.3, a lower value. Furthermore, the voltammograms indicate the presence of two forms, with similar reduction potential. Strangely, *M. nauticus* cytochrome *c*_552_ has a similar pKa observed in the redox potential determined by potentiometry (Saraiva et al., 1994), but not by cyclic voltammetry (data not shown) (Coutinho, 2013).

The observed band at 620 nm decreases in intensity with the increasing of the pH, while the band at 695 nm (due to the axially coordinated methionine) increases in intensity. This can be explained by the fact that the methionine ligand becomes strongly bound to the heme iron, and thus the heme becomes low spin, while at low pHs the methionine can be partially replaced by a low-field ligand, possibly OH^-^, which can be protonated to water. This could explain the observed low-spin EPR signal (Moura et al., 1988), since it can have a temperature-dependent high/low-spin equilibrium (Barreiro et al., 2023a). Another possible explanation for the observed equilibrium could be the protonation/deprotonation of the propionates. In any case both hypotheses do not promote structural changes in the heme pocket since the near-UV/visible CD spectra do not change with pH (Figure S6, in Supplementary Materials).

### 4.3 Thermostability

The thermostability of *W. succinogenes* cytochrome *c*_552_ has been studied for the first time by CD and DSC. The T_M_ values estimated by the two complementary techniques are similar, but no other thermodynamic parameters can be estimated because the unfolding process is irreversible. This *c-*type cytochrome is rather stable, and its stability decreases close to its pI. The effect of pH on the thermostability of some *c*-type cytochromes has been studied at very low pH values (Kuroda et al., 1992). However, there are no reports on the effect of pH on the thermostability of *c*-type cytochromes in the range studied here, so the destabilization at high pH, observed for *Ws* cytochrome *c*_552_, may be due to a change in the hydrogen bonds that stabilize the structure of this protein.

The unfolding temperature observed at pH 7.0 is relatively higher than that observed for other proteins isolated from mesophilic bacteria (Nobrega et al., 2017), although some exceptions have been reported (Barreiro et al., 2023b). However, the T_M_ is relatively low (90 °C, at pH 5.0) compared to that of cytochromes *c*_552_ from thermophilic organisms: *Hydrogenophilus thermoluteolus* and *H. thermophilus*, which have a T_M_ of 108 ° C and 121 °C, respectively, at pH 5.0 (Oikawa et al., 2005; Nakamura et al., 2006). These two *c*-type cytochromes are as small as *Ws* cytochrome *c*_552_, containing about 80 residues. Interestingly, however, although composed of only 4 α-helices, there is a loop protecting the 6-propionate group of the porphyrin ring (Figure S10, in Supplementary Material), which is absent in *W. succinogenes* cytochrome *c*_552_.

Furthermore, *Pseudomonas aeruginosa* cytochrome *c*_551_, being an 82-residue protein from a mesophilic organism, has a T_M_ of 85 °C, at pH 7.0 (Sanbongi et al., 1989), which compares well with the unfolding temperature of *W. succinogenes* cytochrome *c*_552_, at pH 7.0 (80 °C). The structure of this cytochrome *c*_551_ is similar to the previously mentioned highly thermostable *c*-type cytochromes, with the 4 α-helices and the loop protecting the 6-propionate, so perhaps the exposed heme does not affect the thermostability of the protein as much. In fact, some residues have been proposed to be the main cause of thermostability between *H. thermophilus* cytochrome *c*_552_ and *P. aeruginosa* cytochrome *c*_551_ (Oikawa et al., 2005), suggesting that the stabilization may be due to side chain interactions.

## 5 Author Contributions

SRP has planned the experiments with contribution from IM, MSPC and CC. SRP has cloned the cytochrome, CC and NP have performed the cyclic voltammetry, VHM, MSPC and SRP have performed CD and DSC experiments. CC, MSPC and SRP have analyzed the data, with contributions from VHM. SRP has prepared the draft of the manuscript with contributions from the other authors. All authors have read and approved the manuscript.

## 7 Acknowledgments

The authors acknowledge the Biolab for the acquisition of CD and DSC data.

## 8 Supplementary Material

The Supplementary Material for this article can be found online at:

## 9 Data Availability Statement

The authors will provide any of the datasheets presented upon request.

