## SupplementaryFigures for "Effect of pH on the thermostability and redox properties of cytochrome *c*_552_ from *Wolinella succinogenes*"

### Index

**S1. Primary sequence analysis and Alphafold confidence** 2

**S2. PAGE and SDS-PAGE of heterologously produced *Ws* cytochrome $c_{552}$** 2

**S3. Thermogram prior to baseline correction (by DSC)** 3

**S4. CD spectra in the far-UV, near-UV and visible region at different pH values** 3

**S5. Cyclic voltammograms obtained at different pH values** 5

**S6. Structural analysis of *Ws* cytochrome $c_{552}$ and other *c*-type cytochromes** 5

### S1. Primary sequence analysis and Alphafold confidence

The primary sequence of cytochrome $c_{552}$ from *Wolinella succinogenes* was analyzed using SignalP web server to identify the signal peptide.

```
1         |         |         |         |         |        60
MKKIALGMIF AAASLMAADG ATLYKKCVAC HGVKAEKPAL GKSEVIAGWD KAKLVEELKA
YKAGTLNRNG MGAMMKGQMA SFSDADIEAV SEYISTLK
```

**Figure S1.** Primary sequence of cytochrome $c_{552}$ from *W. succinogenes* (WS0700). The sequence peptide that was removed due to cloning is identified in blue. Due to the cloning procedure an additional Met residue was added to the N-terminus.

The mature protein that is heterologously produced was submitted to AlphaFold Colab, and a model structure was obtained, without the heme (Figure S2).

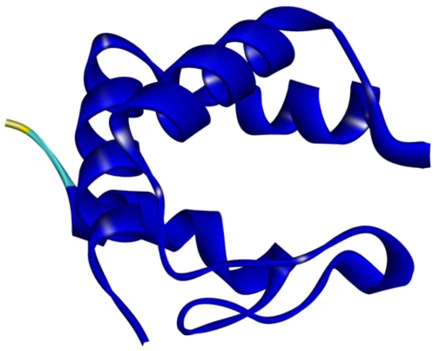

**Figure S2.** Backbone of *Ws* cytochrome $c_{552}$ colored according to per-residue confidence (pLDDT) in a scale from 0-100. Most regions have a pLDDT > 90 (dark blue color), indicating that are expected to be modelled to high accuracy, a short region at the N-terminal (residue Ala2 to Asp3) have a pLDDT between 70 and 90, meaning that they are well modelled, and one residue (Met1) have a low confidence (pLDDT < 50).

### S2. PAGE and SDS-PAGE of heterologously produced *Ws* cytochrome $c_{552}$

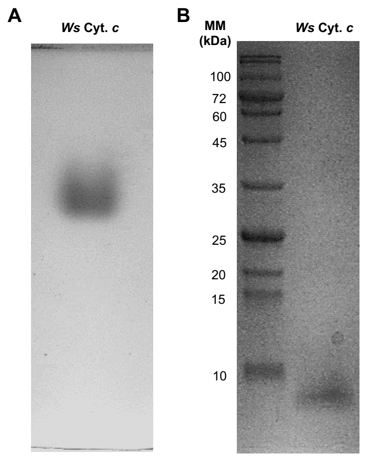


**Figure S3. (A)** 10% PAGE and **(B)** 15% SDS-PAGE of purified *Ws* cytochrome $c_{552}$. PAGE was prepared in Tricine-Imidazol buffer system, while SDS-PAGE was prepared in Tris-Tricine buffer system. Gel stained with Coomassie-blue.

### S3. Thermogram prior to baseline correction (by DSC)

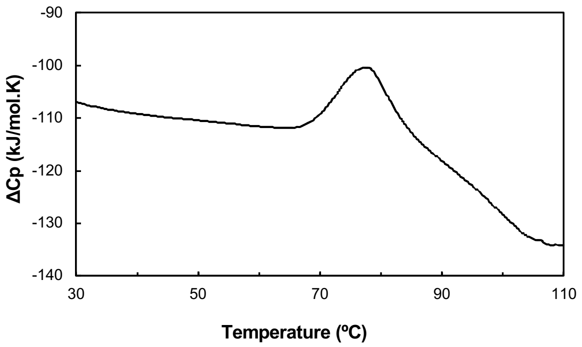


**Figure S4.** Thermogram of 93 μM *Ws* cytochrome $c_{552}$ in phosphate buffer, pH 7.0, prior to baseline correction to show that this protein aggregates in solution upon unfolding. Similar thermogram were observed for the other pHs.

### S4. CD spectra in the far-UV, near-UV and visible region at different pH values

The CD spectra acquired at 25 °C for the different pH values in the far-UV region is shown in Figure S5. Each spectrum has the same data treatment has the one shown in the manuscript (average of three replicates).

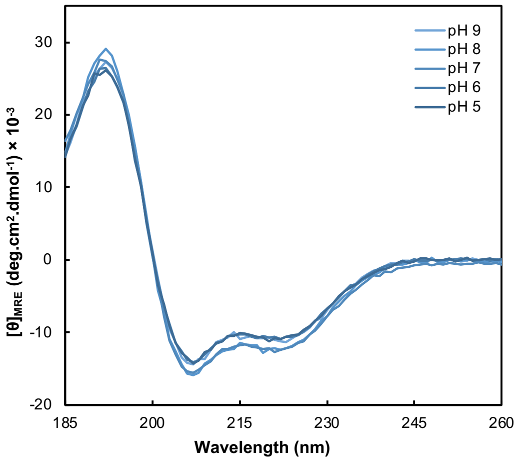


**Figure S5.** CD spectra in the far-UV region of *Ws* cytochrome $c_{552}$ acquired at different pH values, at 25 °C.

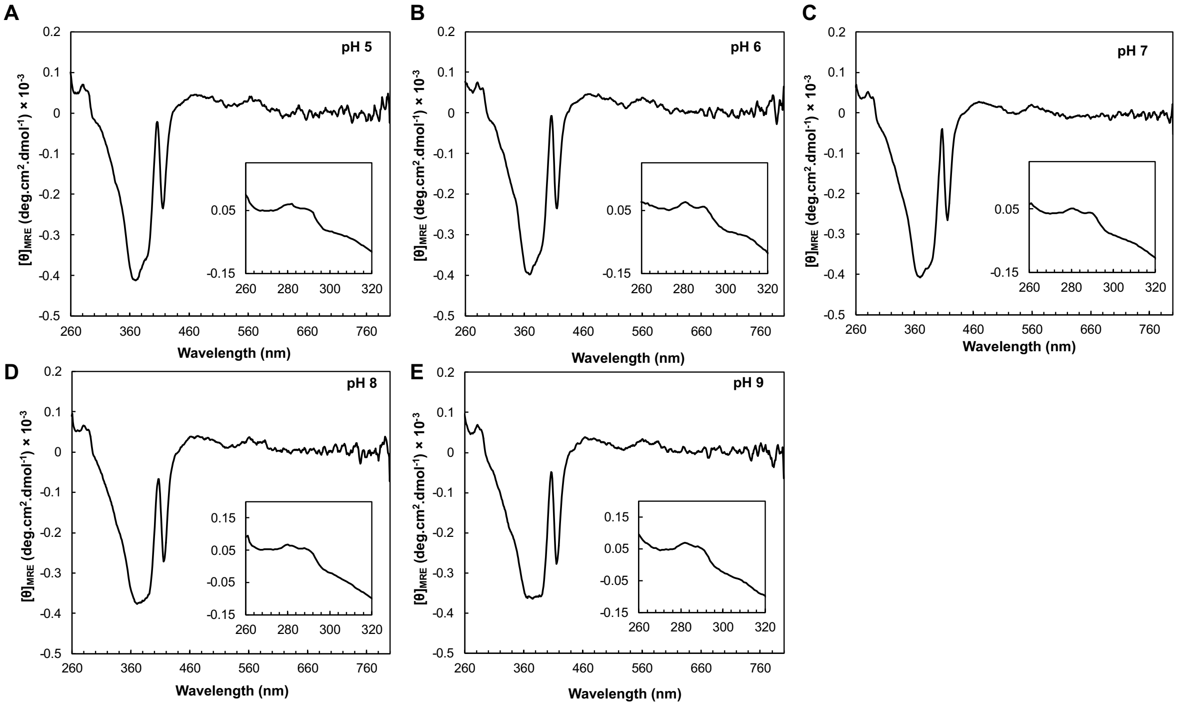


**Figure S6.** CD spectra in the near-UV and visible regions of *Ws* cytochrome $c_{552}$ acquired at different pH values, at 25 °C. (A)

In Figure S7, it is shown the CD spectra prior and after the temperature ramp in the far-UV region for each pH value (Panel A to E), and the profile of the molar ellipticity at 208 nm (Panel F).

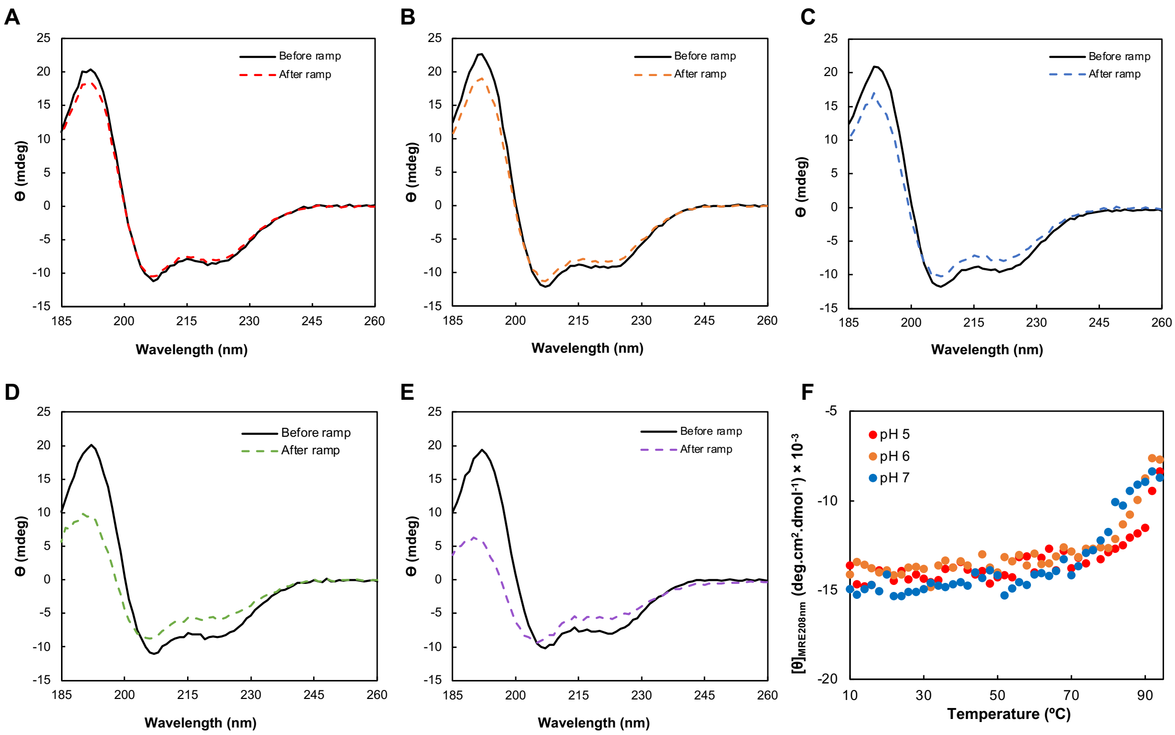


**Figure S7.** CD spectra prior (full line) and after the temperature ramp (dashed line) for each pH value (Panel A to E) and the profile of the molar ellipticity at 208 nm (Panel F) along the temperature ramp, at pH 5-7. Legend: pH 5.0 (red), pH 6.0 (orange), pH 7.0 (blue), pH 8.0 (green) and pH 9.0 (purple). Each spectrum is the average of three, acquired as mentioned in Materials and Methods.

### S5. Cyclic voltammograms obtained at different pH values

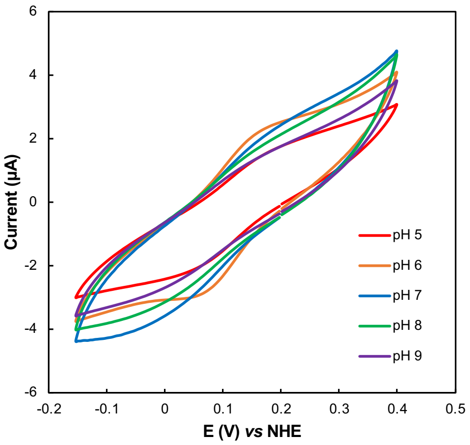


**Figure S8.** Raw data without normalization or baseline corrections. Representative voltammograms of the *Ws* cytochrome $c_{552}$ at different pH values, attained on cysteamine modified gold electrode, at scan rate of 50 mV/s, 20 mM phosphate/acetate buffer, and 0.1 M NaCl, pH 7, room temperature. Data acquired under strict anaerobic environment (anerobic chamber, $O_2$ < 0.1 ppm).

### S6. Structural analysis of *Ws* cytochrome $c_{552}$ and other *c*-type cytochromes

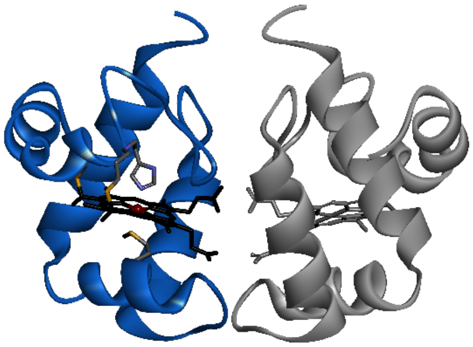

**Figure S9.** Backbone of *M. nauticus* cytochrome $c_{552}$ showing the dimer with the heme propionate groups at the interface. The figure was prepared in BIOVIA Discovery Studio Visualizer with the coordinated 1CNO.

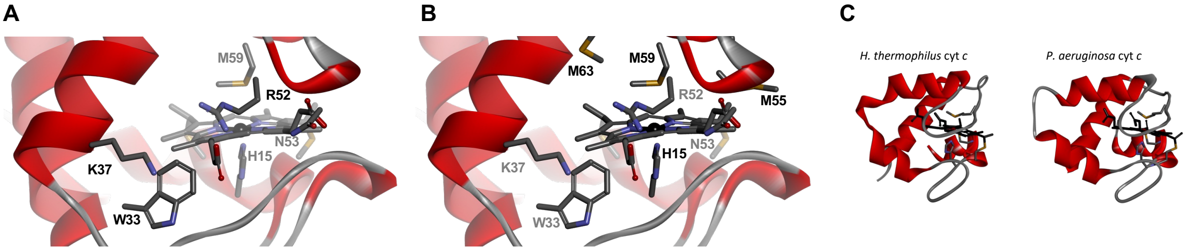


**Figure S10.** Highlights of the model structure of *Ws* cytochrome $c_{552}$, showing the close residues near the propionate groups (Panel A), the several methionine residues close in sequence to the coordinating methionine M59 (Panel B). M58 is outside the figure, solvent exposed. The residues are number according to the mature sequence of the heterologous *Ws* cytochrome $c_{552}$with the additional methionine introduced due to the cloning. In Panel C is shown the structure of *c*-type cytochromes from *P. aeruginosa* and *H. thermophilus* with the loops covering the heme. Figure was prepared in BIOVIA Discovery Studio Visualizer with the coordinates 1AYG and 351C, and the model structure of *Ws* cytochrome $c_{552}$.
